# Transcriptomic profiling implicates PAF1 in both active and repressive immune regulatory networks

**DOI:** 10.1101/2022.03.28.485548

**Authors:** Matthew W. Kenaston, Oanh H. Pham, Marine J. Petit, Priya S. Shah

**Affiliations:** Department of Microbiology and Molecular Genetics, University of California, Davis, Davis, California, USA.; Department of Chemical Engineering, University of California, Davis, Davis, California, USA.

**Keywords:** polymerase associated factor 1 complex (PAF1C), innate immunity, network analysis, gene expression, transcriptomics, regulatory motifs

## Abstract

**Background:** Sitting at the interface of gene expression and host-pathogen interaction, polymerase associated factor 1 complex (PAF1C) is a rising player in the innate immune response. The complex localizes to the nucleus and associates with chromatin to modulate RNA polymerase II (RNAPII) elongation of gene transcripts. Performing this function at both proximal and distal regulatory elements, PAF1C interacts with many host factors across such sites, along with several microbial proteins during infection. Therefore, translating the ubiquity of PAF1C into specific impacts on immune gene expression remains especially relevant.

**Results:** Advancing past work, we treat PAF1 knockout cells with a slate of immune stimuli to identify key trends in PAF1-dependent gene expression with broad analytical depth. From our transcriptomic data, we confirm PAF1 is an activator of traditional immune response pathways as well as other cellular pathways correlated with pathogen defense. With this model, we employ computational approaches to refine how PAF1 may contribute to both gene activation and suppression. Specifically focusing on transcriptional motifs, we predict gene regulatory elements strongly associated with PAF1, including those implicated in an immune response. Overall, our results suggest PAF1 is potentially involved in innate immunity at several distinct axes of regulation.

**Conclusions:** By identifying PAF1-dependent gene expression across several pathogenic contexts, we confirm PAF1C to be a key mediator of innate immunity. Combining these transcriptomic profiles with potential regulatory networks corroborates the previously identified functions of PAF1C. With this, we foster new avenues for its study as a regulator of innate immunity, and our results will serve as a basis for targeted study of PAF1C in future validation studies.

## Background

A robust innate immune response relies on the early detection of pathogen-associated molecular patterns (PAMPs) by host pattern recognition receptors. This sensing triggers various signaling pathways culminating in responsive gene expression meant to prevent or combat infection. The polymerase associated factor 1 complex (PAF1C) is increasingly implicated in such a response at the level of gene expression—resulting from its key roles in transcriptional elongation, chromatin remodeling, and histone modification. The primary members of this nuclear complex are PAF1, LEO1, CTR9 and CDC73 [1]. First discovered as a homolog in yeast [2], PAF1C serves as a scaffold for recruitment of histone- and RNA polymerase II (RNAPII)-modifying enzymes [3–7]. At these sites, PAF1C activity regulates the release of a paused RNAPII at promoters, permitting productive elongation of the transcript [8–10].

Interestingly, PAF1C occupation of certain super-enhancers in cancer cells results in negative regulation of associated genes [11]. As such, loss of PAF1C in certain contexts can activate gene expression. Other work has attributed enhancer occupation to PAF1C’s modulation of RNAPII elongation rates rather than direct suppression [12]. Though the exact mechanisms behind these divergent functions are not fully understood, PAF1C is poised to extensively modulate gene expression depending on the genes and regulatory elements present.

PAF1C regulation of gene expression impacts the ability of cells to mount an antimicrobial response. Specifically, PAF1 depletion has been shown to disrupt interferon-related signaling and lead to increases in viral replication for both dengue virus and influenza A virus (IAV) [13–15]. These same studies found such viruses target PAF1 to abrogate its activation of an immune response, and, recently, similar antagonism has been discovered in bacteria [16]. PAF1 and various complex members also act as restriction factors for HIV-1 [17–19]. Others have shown PAF1C to increase tumor necrosis factor (TNF) expression as part of the inflammatory response [20]. Thus, PAF1C is emerging as an influential factor in maintaining host innate immunity, though its scope of regulation remains imprecise.

Here, we computationally interrogate the extent and conservation of PAF1C-mediated gene expression by treating PAF1 knockout (PAF1 KO) cells with a diverse set of immune stimuli, including well-recognized PAMPs. Building on previous work, we confirm PAF1 promotes an antiviral response. We further identify the complex as not only a modulator of innate immunity, but a suppressor of mechanisms that induce a proviral state in cells. By identifying a conserved set of genes that are PAF1-dependent, we construct various models to predict potential transcription factors and regulatory elements that may interact with PAF1C to activate various innate immune pathways. Ultimately, our results strengthen the model of PAF1C as a crucial player in the host immune response while providing the necessary foundation for future work.

## Results

### Shared and distinct immune response pathways in A549 cells

To broaden our identification of PAF1-dependent gene expression, we sought to activate multiple immune pathways. This way, we could differentiate a core set of genes correlated to PAF1 while preserving any nuance for a particular stress response (Table 1). Based on our work using a dsRNA mimic in PAF1 KO cells, we hypothesized that genes both upstream and downstream of type I interferon (IFN-I) signaling were PAF1-dependent [14]. PAF1 KO and non-targeting control (ncgRNA) cells were previously developed in A549 lung epithelial cells, which have been used extensively for the study of the innate immune response [21–23]. We have shown PAF1 Rescue cells recapitulate PAF1-dependent gene expression [14]. Therefore, we opted to use ncgRNA cells as a universal control that has the potential to enable comparisons across multiple gene knockouts in the future.

**Table 1.**
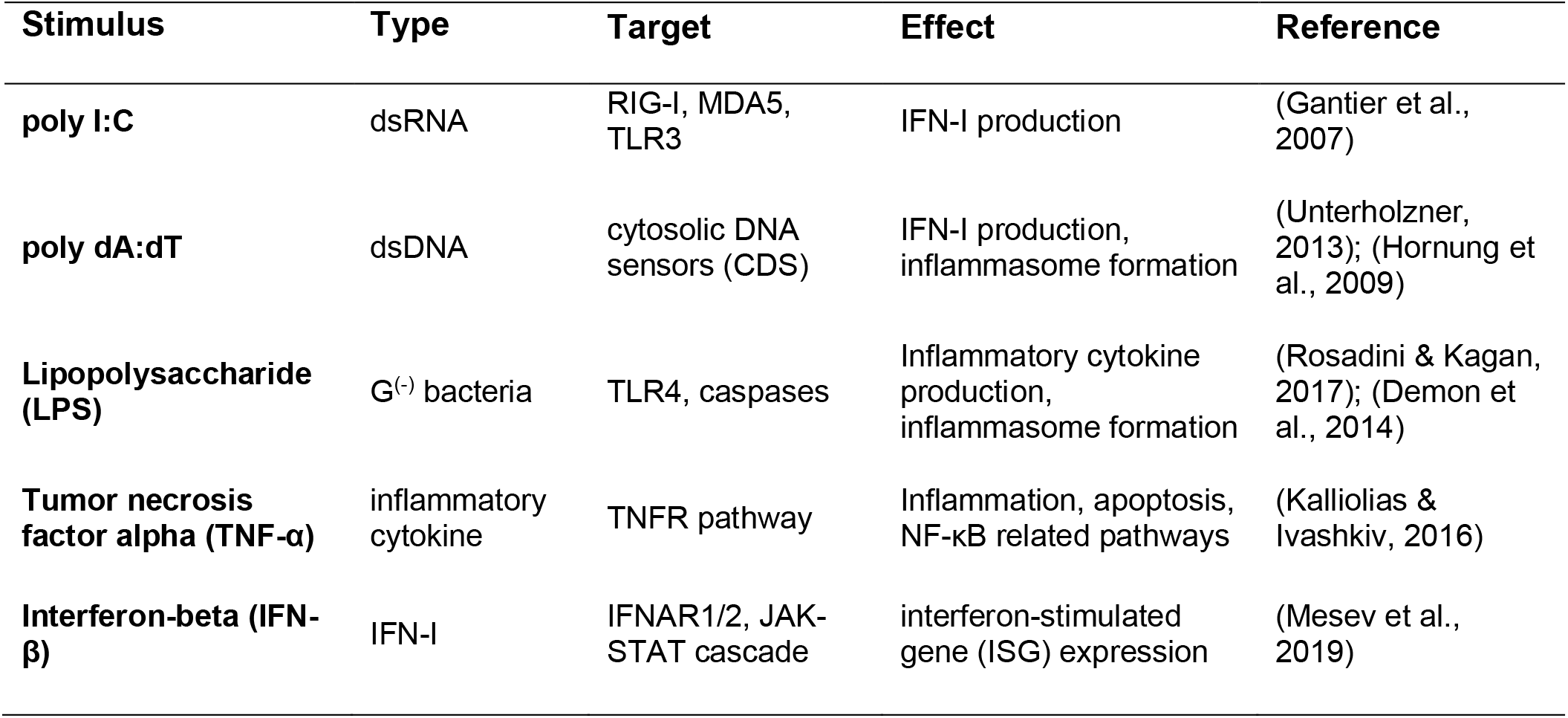
Summary of innate immune stimuli.

Both cell lines were treated with the selected stimuli for three hours and subjected to 3’ Tag-sequencing (Fig. 1A). Though PAF1 nuclear localization may be marginally disrupted in A549 cells under poly I:C-induced stress, this trend was not observed for LPS or IFN-β (Fig. S1). To capture all the variability of our processed reads, cell genotype (PAF1 KO, ncgRNA) and immune stimuli were treated as distinct conditions within the computational design of our differential gene expression analysis. This allowed for numerous direct comparisons to be made while accounting for inherent differences amongst those factors.

**Figure 1.**
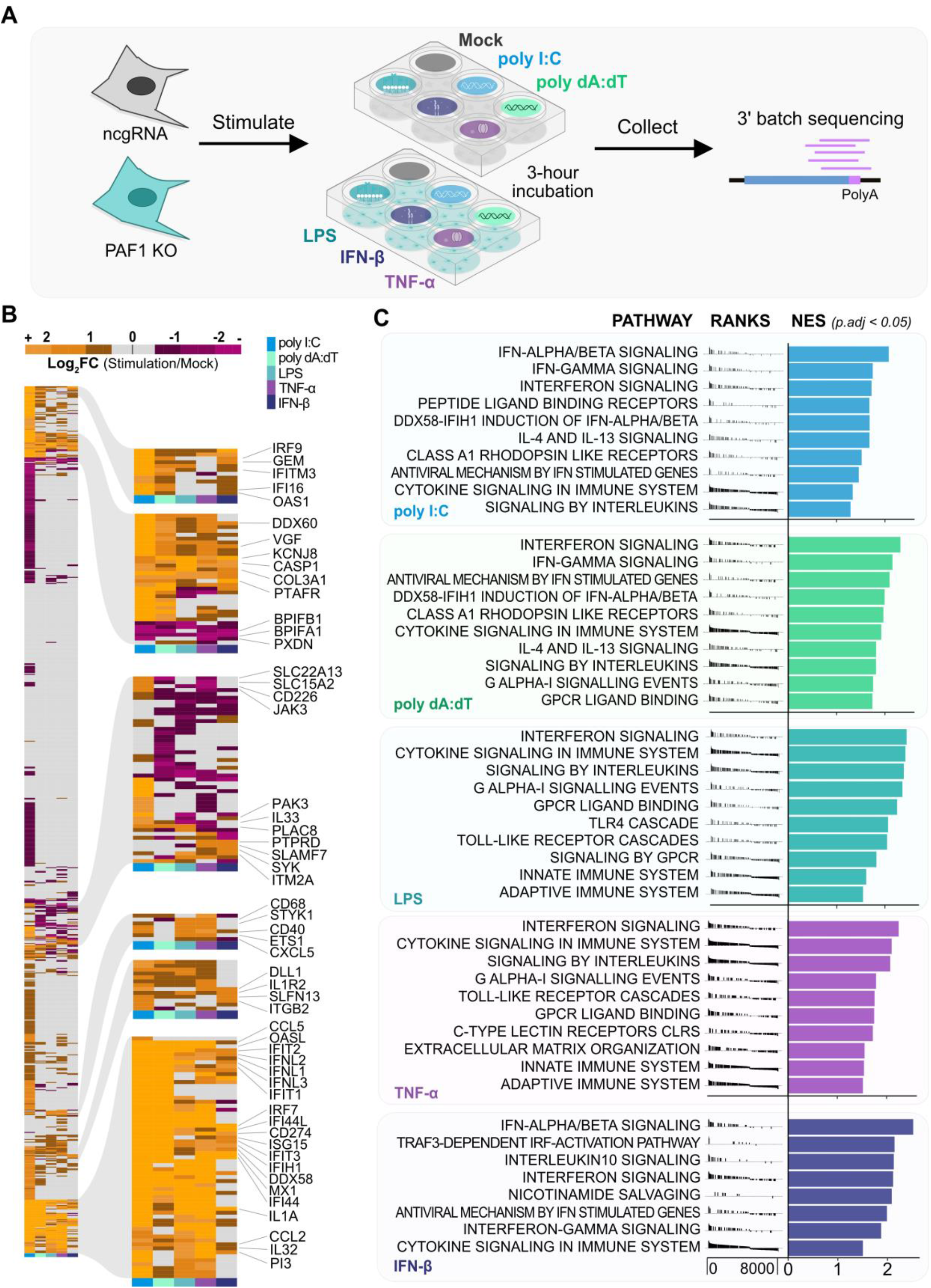
Stimuli activate shared and distinct pathways of the innate immune response. (A) Across 3 biological replicates, ncgRNA (grey) and PAF1 KO (teal) cells were subject to a 3-hour stimulation by the treatments described. Following, RNA was extracted, purified, and sequenced. Aligned reads underwent DESeq2 differential gene expression analysis. (B) A heatmap of log2 foldchanges (ncgRNA stimulation / ncgRNA mock) for immune response genes (GO:0002376) confirms immune activation. Hierarchal clustering using Ward’s method identified notable clusters of genes for labeling. (C) GSEA was performed on DEGs in ncgRNA cells for each stimulation, wherein genes were ranked by a weighted statistic for log2 foldchange and adjusted p-value. The normalized enrichment scores (NES) for, at most, the top 10 enriched Reactome pathways were plotted for each stimulus (p.adj < 0.05). The full GSEA output is available in Table S2A.

We first analyzed the data by focusing on the ncgRNA control cells treated with various stimuli to confirm that these treatments induced immune response gene expression compared to mock-treated cells. We found each stimulus activated typical genes associated with the immune response (GO:0002376) (Fig. 1B, Table S1A). The inflammatory stimuli (TNF-α, LPS) appeared to activate some genes distinct from the interferon-related stimuli (IFN-β, poly dA:dT, poly I:C), and vice versa. These results align with principal component analysis (PCA), which shows two axes of distinct immune activation (Fig. S2A). Poly I:C was the strongest stimulus as measured by number of genes induced. The same trends are reiterated upon identifying the top differentially expressed genes (DEGs) (|log2 fold-change| > 1.5, p-value < 0.05) for each stimulus (Fig. S2B) and validating specific gene expression by qRT-PCR (Fig. S2C).

We next implemented gene set enrichment analysis (GSEA) to measure induction of the immune response in an unbiased manner, using Reactome gene sets encompassing a broad number of cellular pathways. With this, we confirmed all stimuli activated known immune response pathways (Fig. 1C). Corroborating previous observations, our inflammatory stimuli induced some unique pathways, as did the pathways activating the interferon response. Specifically, broad GSEA revealed each stimulus as a potent driver of interferon-related pathways, including interferon alpha-beta and gamma (Fig. 1C). Given their mediation of the inflammatory response, TNF-α and LPS were more enriched for pathways related to the adaptive immune response and TLR signaling cascades compared to their counterparts (Fig. 1C, Table S2A). Overall, our stimuli are broad and effective inducers of the immune response.

### PAF1 KO results in dysregulation of immune gene expression

To discern whether PAF1-mediated gene expression is differentiated across immune stimuli, we compared PAF1 KO cells to the control ncgRNA cells for every stimulus. We performed PCA on immune genes while excluding batch effects and variance attributed to differences previously found amongst the stimuli. In doing so, we found immune gene expression in PAF1 KO cells is distinct from that of ncgRNA cells, accounting for 72% of the remaining variance (Fig. 2A).

**Figure 2.**
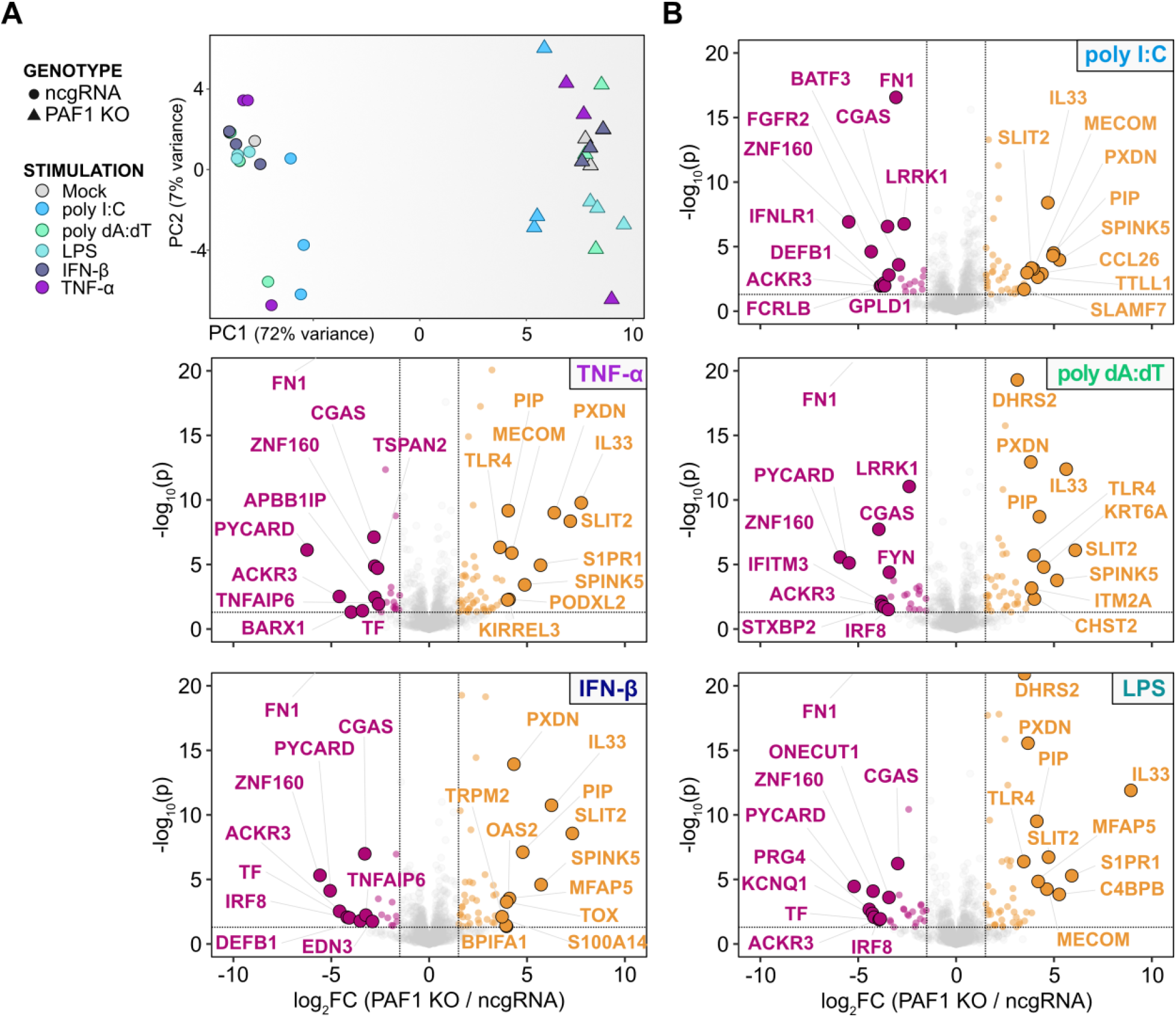
PAF1 KO cells exhibit a distinct immune gene expression profile. (A) PCA shows distinct gene expression profile of PAF1 KO relative to ncgRNA. The counts for immune genes (GO:0002376) were corrected with a variance-stabilizing transformation via DESeq2, and the variability of biological replicates and immune activation were removed as batch effects with ComBat. PCA was performed on the resulting counts, with the top 2 principal components being plotted. Cell lines are represented by shape (circle: ncgRNA, triangle: PAF1 KO) and stimuli are dictated by color. (B) Volcano plots show differential gene expression of immune genes in PAF1 KO relative to ncgRNA across stimuli. Contrasts were performed within the DESeq2 pipeline. The resulting log2 foldchanges and –log10 p-values are plotted for immune genes, with the top-10 positive and negative log2 foldchanges being highlighted/labeled. Cutoffs are set at log2 foldchange > 1.5 or < −1.5, and p-value < 0.05.

For each comparison of PAF1 KO to ncgRNA, we identified the top immune system DEGs (GO:0002376, |log2 foldchange| > 1.5, p-value < 0.05). Across stimuli, we found several immune genes which are both upregulated and downregulated in PAF1 KO relative to ncgRNA (Fig. 2B, Fig. S3). cGAS, a well-recognized cytosolic DNA sensor [24], had significantly lower expression in PAF1 KO cells compared to ncgRNA cells for all stimuli (Fig. 2B, Fig. S3, Table S1B). We found similar trends for the expression of PYCARD/ASC, a key activator of inflammatory caspases for subsequent maturation of IL-1β [25] and mediator of NF-κB related apoptosis [26]. The same was also found for IRF8 and LRRK1, both of which are regulators of NF-κB activation for various immune pathways [27, 28].

Similar to previously published studies indicating a negative regulatory role for PAF1, we also identified genes whose expression increased following PAF1 KO. In general, these genes were more variable in terms of immune modulation. Several of them are known to repress the immune response under various cellular or pathogenic contexts—including IL33, S1PR1, and MECOM [29–33](Fig. 2B, Fig. S3). However, other upregulated genes, such as OAS2, PXDN, and TLR4, are predominantly mediators of immune activation and antimicrobial defense [34–36]. TLR4 gene expression is also suppressed by ZNF160 [37], a gene with consistent downregulation in PAF1 KO (Fig. 2B, Table S1B). Thus, some of these effects may be secondary to a direct effect by PAF1 on the gene of interest. Overall, the apparent upregulation and downregulation of gene expression following PAF1 KO further reinforces the model of PAF1 suppressing some gene expression mechanisms while likely inducing others.

### Immune pathway analysis of PAF1-dependent gene expression

Next, we applied an increasingly targeted approach to identify how altered gene expression may modulate the innate immune response. First, we chose to ascertain whether immune induction in PAF1 KO was reduced compared to ncgRNA control cells. When comparing stimuli against the mock-treatment for both cell lines, we found an overall decrease in immune gene expression for PAF1 KO (Fig. 3A). Though these changes were relatively modest, they were consistent and resulted in strong statistical significance for each comparison. Importantly, some genes happened to be more induced in PAF1 KO (Fig. 3A), which aligns with our previous analysis of individual DEGs (Fig. 2B). Thus, PAF1 is evidently required for robust immune gene expression regardless of downstream effect, yet likely fills suppressive roles as well.

**Figure 3.**
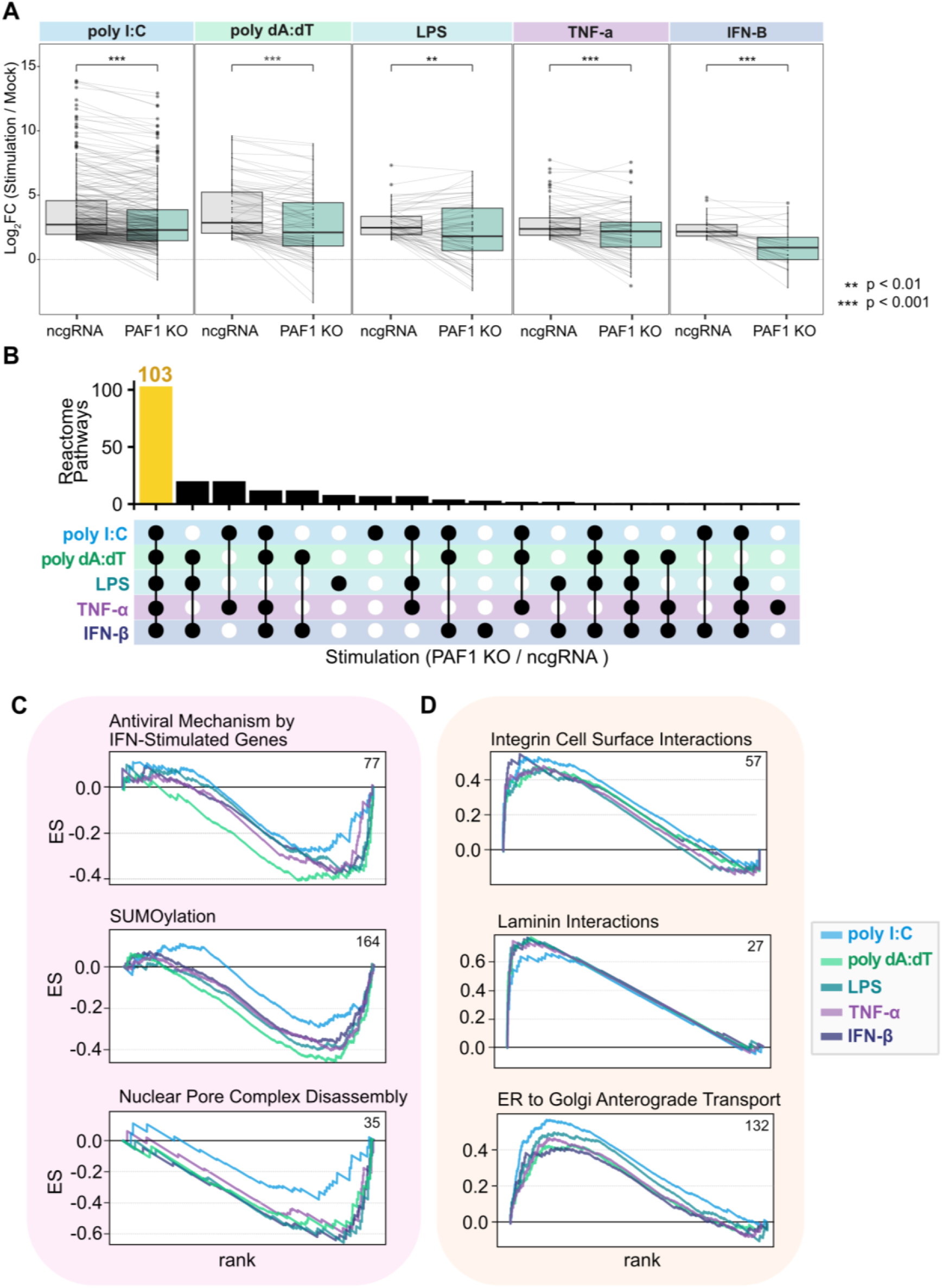
PAF1 modulates the expression of genes related to immune mechanisms and host machinery. (A) The log2 foldchanges of immune genes (GO: 0002376) upregulated (log2 foldchange > 2) following each stimulus in ncgRNA cells (relative to the mock) are plotted. These genes are mapped to their counterpart log2 foldchanges in PAF1 KO cells. A paired Wilcoxon signed rank test with Bonferroni correction was performed to assess significant changes between the distributions of foldchanges. (B) GSEA was performed on DEGs from PAF1 KO (relative to ncgRNA) for each stimulus, wherein genes were ranked by a weighted statistic for log2 foldchange and adjusted p-value. Significantly enriched Reactome pathways (p.adj < 0.05) were analyzed for overlap amongst stimuli and the results shown on an upset plot. The number of pathways that overlap across all stimuli is highlighted in yellow. Filled dots indicate which stimuli shared enrichment for the given number of pathways. (C-D) GSEA random walks with the running total enrichment score (ES) were plotted for Reactome pathways found to be negatively (C) or positively (D) enriched across stimuli. Colors differentiate stimuli on each plot. Numbers listed in the upper-right corner of plots indicate the size of the gene set shown. The full GSEA output is available in Table S2B.

Seeking to validate this result, we again used GSEA to identify pathways that are PAF1-dependent. We performed GSEA on our DEGs, evaluating pathway overlap to ascertain which phenotypes may be conserved or differentiated across stimuli. Most significantly enriched pathways (p.adj < 0.05) were shared across stimuli, totaling to 103 pathways (Fig. 3B). We hypothesized that an antimicrobial effect of PAF1 could derive from positively regulated genes that are antimicrobial, or negatively regulated genes that are promicrobial. Therefore, we investigated these pathways for those that were downregulated and antimicrobial, or upregulated and promicrobial. Conserved downregulated pathways included the antiviral mechanism by IFN-stimulated genes as well as SUMOylation (Fig. 3C). Both pathways are predisposed to contribute to a defensive state in cells if active [38–42]. We also found negative enrichment of pathways associated with the nuclear pore complex (Table S2B), including disassembly (Fig. 3C). Therefore, PAF1 may facilitate innate immune defense beyond traditional pathways alone.

Expanding on the potentially suppressive role of PAF1, we also identified upregulated pathways in PAF1 KO. Strong positive enrichment is evident for both laminin and integrin cell surface interactions (Fig. 3D, Table S2B), both correlated with sites prone to pathogen adhesion and entry [43–47]. Similarly, ER to Golgi anterograde transport was also positively enriched (Fig. 3D), a process coopted by some viruses for egress [48]. This suggests PAF1 activates certain defensive mechanisms in tandem with suppressing pathways prone to microbial hijacking.

To see if this trend could be validated for specific viruses, we next considered whether PAF1 is suppressive of viral host dependency factors, as shown with flaviviruses previously [14]. Across stimuli, GSEA revealed that *Flavivirus* host-dependency factors were induced following PAF1 KO (Fig. 4A-B), suggesting PAF1 may be suppressive of those factors typically. For Influenza A virus (IAV), this trend was present yet reduced across stimuli (Fig. 4A, C). However, the Reactome pathway for NS1-mediated effects on host pathways was significantly downregulated for all stimuli in PAF1 KO (Table S2B), which is expected because NS1 targets PAF1 for antagonism [13]. We also assessed host dependency factors for HIV and SARS-CoV-2 given their implication as targets of PAF1-mediated restriction [19, 49]. Like IAV, there was a strong effect under poly I:C stimulation for these viruses, but not the other stimuli (Fig. 4A, D-E). Hence, the extent of PAF1 restriction by suppression of dependency factors may depend on the type and degree of immune activation in the first place.

**Figure 4.**
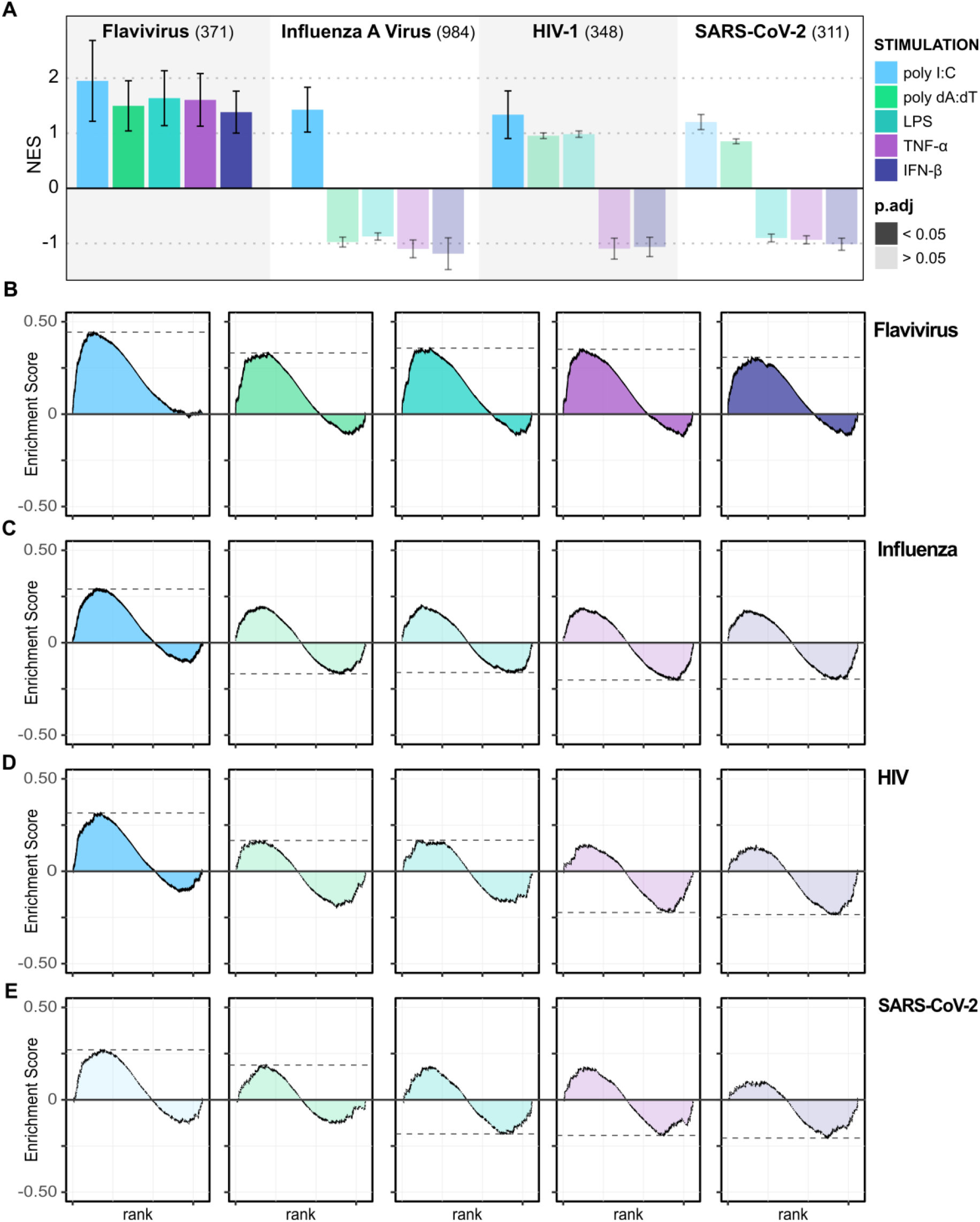
PAF1 KO shows increased expression of virus host-dependency factors. GSEA was performed on DEGs in PAF1 KO relative to ncgRNA for each stimulation using custom gene sets describing host-dependency factors for various viruses. Colors indicate stimuli type and transparency indicates adjusted p-values for each test. (A) NES values are plotted with error bars representing log2 of the expected error for the standard deviation of the P-value logarithm. The size of each gene set is provided in parentheses. GSEA random walks as summarized in (A) are shown for *Flaviviruses* (B), Influenza A Virus (C), HIV (D), and SARS-CoV-2 (E).

Considering these deviations in PAF1-dependency across stimuli, we hypothesized there could be stimulus-specific gene expression that was also PAF1-dependent. Many of these stimulus-specific genes were attributed to different degrees of immune activation relative to the mock. Yet, we still identified some immune genes which were consistently activated across stimuli in ncgRNA cells but were stimulus dependent in PAF1 KO (Fig. S4A-B). For example, REC8 was significantly downregulated under nucleic acid mimic stimuli (poly I:C, poly dA:dT) in PAF1 KO cells, but not for inflammatory stimuli (LPS, TNF-α) (Fig. S4A). Interestingly, REC8 promotes innate immune signaling via stabilization of MAVS and STING [50], connecting back to our previously identified dysregulation of the cGAS-STING pathway (Fig. 2B). We also compared upstream PAMPs (poly I:C, poly dA:dT, LPS) to downstream activators (TNF-α, IFN-β), but found few genes strongly induced across stimuli in ncgRNA cells, yet differentially expressed across stimuli in PAF1 KO (Fig. S4B). Moreover, none were strongly associated with an immune response. Thus, though PAF1C can function in a stimulus-dependent manner [51–53], this does not appear to be the central driving factor for innate immune modulation.

### Predicted transcriptional motifs underlying PAF1-dependent immune regulation

Since most of the holistic trends we identified were shared across stimuli, we sought to understand how this may translate to consistency across PAF1-dependent expression for individual genes. To that end, we queried all significant DEGs (p.adj < 0.05) from our individual comparisons and aggregated them to identify overlaps. There were also numerous genes associated with only some stimuli but not others, reinforcing that other inputs also contribute to the expression of PAF1-dependent genes. Despite this variability, we discovered 50 downregulated genes and 104 upregulated genes shared across all stimuli (Fig. 5A).

**Figure 5.**
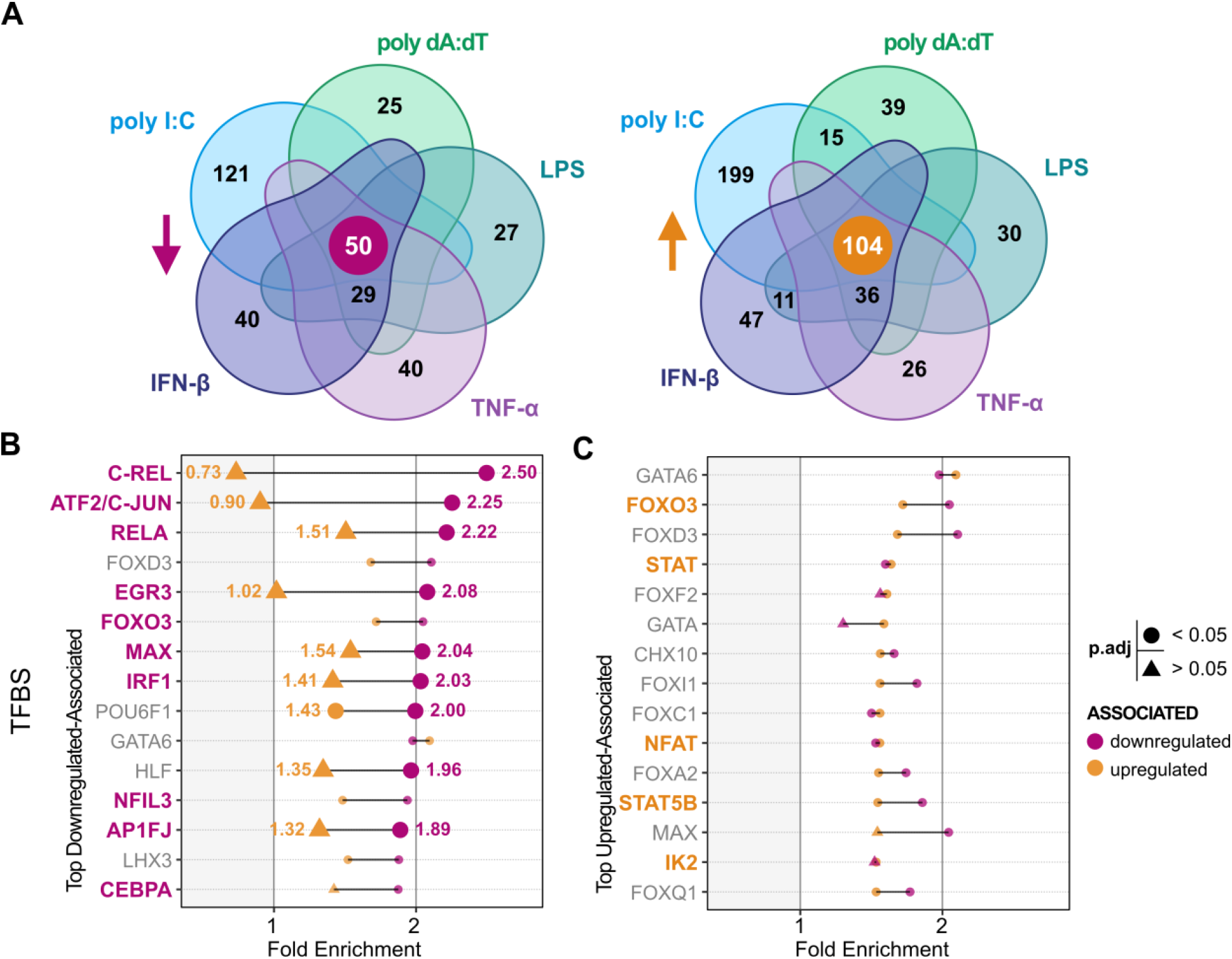
Promoters of conserved PAF1-dependent genes are associated with immune-related transcription factors. (A) Venn diagrams show overlapping DEGs (p.adj < 0.05) from comparison of PAF1 KO versus ncgRNA for each stimulus. The diagram on the left shows downregulated genes and the one to the right shows upregulated genes. Overlaps having greater than 10 genes were labeled. (B-C) Core genes (marked at the center of Venn diagrams) were subjected to TFBS enrichment analysis with DAVID. The fold enrichment (FE) of the top 15 associated TFBS for downregulated (B) and upregulated (C) genes are shown. Emphasis (larger size and labeling) was given to those with greater than 0.5 difference in FE compared to the opposing gene list (downregulated versus upregulated (B), and vice versa (C)). Statistical significance for enrichment was calculated with the Bonferroni method and is shown by shape (circle: p.adj < 0.05, triangle: p.adj > 0.05). TFBS associated with the immune response and significantly enriched have bolded and colored labels.

We performed gene ontology (GO) enrichment analyses on these core genes, but we did not identify any sets that were significantly enriched. PAF1C is known to interact with several transcription factors during the recruitment of RNAPII [7, 51, 54, 55]. Thus, we extended our approach to focus on regulatory elements associated with core genes. Taking the downregulated genes conserved across stimuli, we identified significant enrichment for specific transcription factor binding sites (TFBS) associated with the promoters of those genes (Fig. S5A). Many of the top hits were implicated in the innate immune response, including c-Rel, ATF2/c-Jun, RELA/NF-κB p65, IRF1, AP-1, and EGR3 (Fig. 5B, Table S3A). Supporting the model in which PAF1 is an activator at these sites, there was comparatively less enrichment amongst upregulated genes for these same TFBS. Specifically, c-Rel, ATF2/c-Jun, and EGR3 all displayed greater than 2-fold enrichment for downregulated genes. However, their upregulated-associated counterparts were approximately less than or equal to 1-fold enrichment (no enrichment) (Fig. 5B, Table S3A). This same analytical approach was performed to identify the top TFBS associations for upregulated genes. Yet, this resulted in enrichments of smaller magnitude and less often associated with immune-related TFBS (Fig. 5C). What’s more, these top associations were often outperformed by their downregulated-associated counterparts. Taken together, PAF1-dependent genes have promoters associated with known transcription factors, including those involved with regulation of innate immunity.

We next investigated enhancers associated with our genes because PAF1 could occupy these sites for activation or suppression [8, 9, 11, 12]. We extracted all the genomic regions of enhancers associated with our core genes (Fig. S5A, Table S5B). Mapping the identified sites onto the human genome, we evaluated if PAF1-associated enhancer occupancy was distinct between upregulated and downregulated genes. Strikingly, upregulated and downregulated genes shared little overlap between their respective enhancers (Fig. 6A). When this was tested with a Jaccard statistic, we could only find a single region of overlap, resulting in an extremely low score of 5.55 x 10^-5^. Our downregulated genes had enhancers that were more widely distributed than the upregulated genes (Fig. S5B). Additionally, for the number of genes in between the enhancer and the gene of interest, upregulated-associated enhancers were found to be closer on average (two-tailed T-test, p value < 0.001) (Fig. S5C).

**Figure 6.**
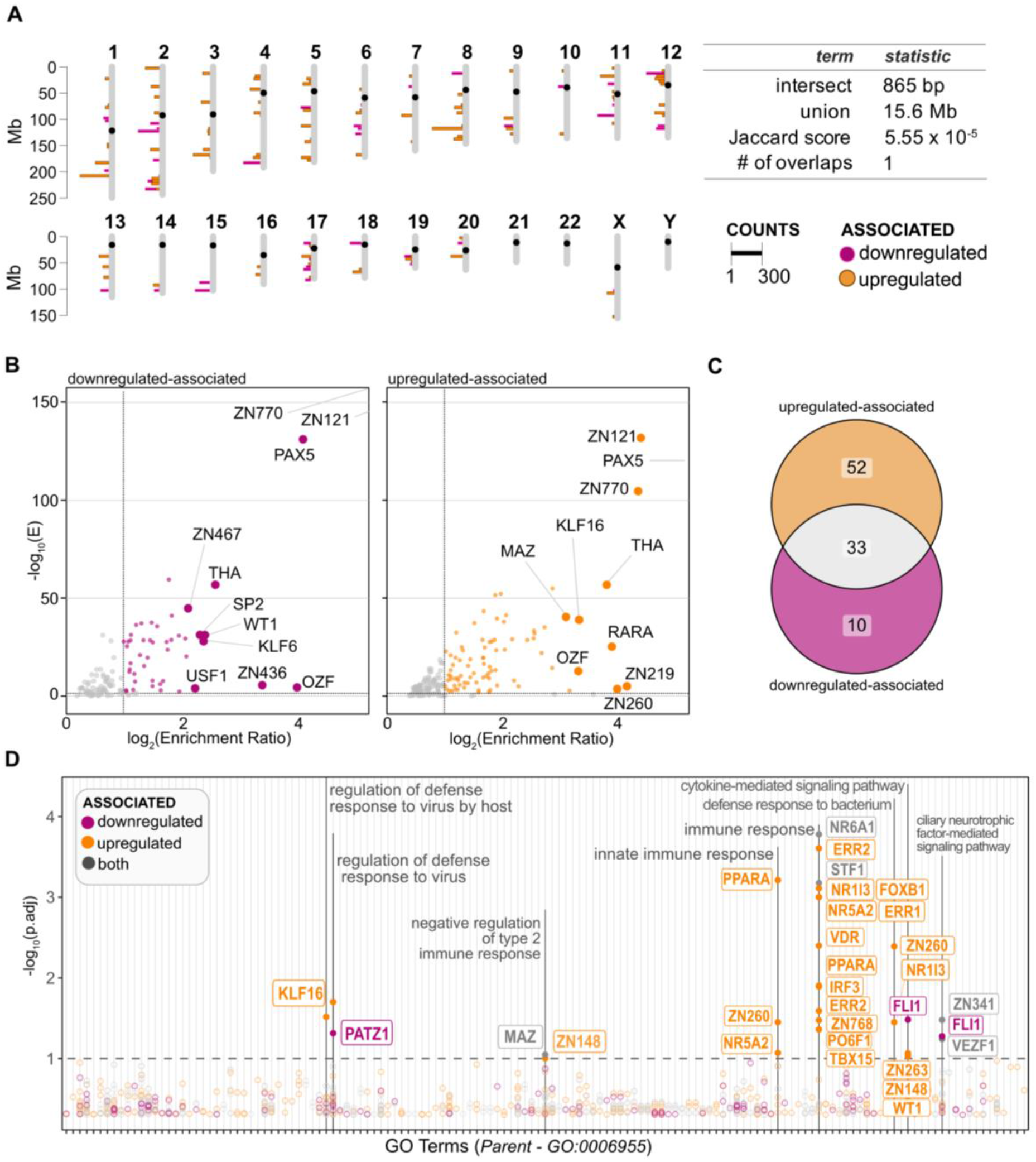
Predicted enhancers associated with PAF1-dependency and the innate immune response. (A) Enhancers associated with the core PAF1-dependent upregulated (orange) and down-regulated (magenta) genes are displayed on the human genome. Enhancer chromosomal regions were extracted and plotted in 0.5 Mb bins. The extent these regions overlapped was calculated with a Jaccard statistic. (B) Volcano plots show results of relative enrichment analysis on extracted enhancers. Significantly enriched motifs (E < 0.05, enrichment ratio > 2) are colored. The top 10 enrichment ratios for known motifs are highlighted for each set. (C) Venn diagram shows overlap of enriched motifs associated with upregulated and downregulated genes. (D) Manhattan plot displays significant (p.adj < 0.05) motifs associated with GO-BP immune response terms (GO: 0006955). Motifs were submitted to GOMo and enriched for correlated terms, correcting for false-discovery rate with Benjamini-Hochberg method. Color indicates whether the enriched motif was associated with downregulated (magenta) or upregulated genes (orange) alone, or if it was shared between them (grey). The analytical pipeline for these analyses is further clarified in Figure S3A.

Although the genomic regions differed, we also considered whether motifs in these enhancers differed as well. For both upregulated and downregulated genes, we discovered several significantly enriched motifs (p.adj < 0.05, log2 fold enrichment > 1) (Fig. 6B-C, Table S3C). Primarily, top motifs for both groups were often assorted zinc-finger binding motifs or those associated with thyroid hormone receptor-related factors (Fig. 6B). Approximately one third of these motifs were shared between upregulated and downregulated genes (Fig. 6C), suggesting these motifs are not necessarily driving the direction of PAF1 regulation for upregulated and downregulated genes. Importantly, there were unique enhancer motifs for both gene sets, so PAF1’s differential gene regulation may be due to these motifs.

To extend this predictive model, we evaluated all significantly enriched motifs for enrichment of GO biological processes (GO-BP) related to the innate immune response (GO: 0006955) (Fig. S5A). Implementing GOMo (Gene Ontology for Motifs), we evaluated whether potential gene targets associated with our enhancer regions were enriched for GO-BPs. We discovered a subset of our motifs were significantly enriched (p.adj < 0.05) for immune-related terms, including innate immunity and the defense response to virus (Fig. 6D, S3D Table). Contrary to our TFBS promoter analysis (Fig. 5B), the enhancers associated with upregulated genes comprised most of the motifs related to the immune response (Fig. 6D). Although motifs associated with downregulated genes or with both sets were less often enriched for GO-BPs, a large fraction still had known immune gene targets (Fig. S6). This illustrates how PAF1 activity at different regulatory elements could be highly variable in regard to innate immunity.

## Discussion

Expanding the cellular role of PAF1C, here we explore the extent of PAF1-dependent gene expression as it relates to the immune response. Not only do we reinforce results from previous studies, but we delineate potential mechanisms that determine and contribute to the degree and type of PAF1-mediation. We show that PAF1 promotes gene expression across a broad range of genes in response to diverse immune stimuli. We also provide additional support that PAF1 can negatively regulate some genes, including proviral host factors. We show these trends in gene expression correlate with known immune response motifs at the promoter and enhancer level. With this work, we refine the role of PAF1C in responding to microbial stress and identify potential immune regulatory networks to be investigated in subsequent studies.

Our work also contributes to reconciling the recently described, divergent functions of PAF1C. From one perspective, PAF1C allows the proximal release of a paused RNAPII from genes, resulting in activation [8]. Yet, the complex can also be a negative regulator of this process [9], maintaining the paused status of RNAPII, especially at super-enhancers in cancer cell lines [11]. Recent work has combined these models by suggesting loss of PAF1C changes the distribution and rates of RNAPII transcription [12], delineating between proximal and distal genomic sites. Building on these models, we show that PAF1 activation is indeed correlated with promoter elements while suppression is more often associated with enhancer sites. Experimental validation of these trends in future work will be pivotal to fully integrating the two models of PAF1C.

At the level of immune gene expression, PAF1C-mediated activation is apparent and aligns with past work. The diminished expression of cGAS and PYCARD/ASC across all stimuli (Fig. 2B), for example, mirrors our group’s past work in finding cGAS expression to be reduced following PAF1 knockout in poly I:C-stimulated cells [14]. Considering both cGAS and ASC are involved in dsDNA-mediated immune activation and exhibit cross-regulation [56], it is particularly interesting to see their shared downregulation. Also, we detected consistently increased expression of IFI16 in PAF1 KO (Table S1B), which can function cooperatively with cGAS [57], suggesting its overexpression may reflect additional dysregulation of this system. Given the cGAS-STING pathway and formation of the AIM2 inflammasome are activated by diverse pathogens regardless of dsDNA genome status [58, 59], PAF1 mediation of their expression could be protective against cellular vulnerability. This is confirmed when PAF1 KO displays a loss in gene expression for the antiviral mechanism by IFN-stimulated genes (Fig. 3).

Though loss of canonical pathways for immune defense is evident, we found additional cellular pathways significantly affected by PAF1 and still linked to immune defense, though less directly. Decreased expression of SUMOylation pathways in our PAF1 KO cells (Fig. 3B) is especially interesting considering all PAF1C members are a target of SUMO to activate the interferon response and counteract IAV [60]. This suggests PAF1C may rely on a potential positive feedback mechanism to increase its mediation of an antimicrobial response. We also highlight decreased expression of genes related to nuclear pore complex maintenance, for their expression has been correlated with immune effector import into the nucleus [61–63]. For this reason, nuclear pore complexes are often targeted by various viruses for degradation [64–68]. Since PAF1 likely facilitates immune gene expression reliant on the translocation of cytoplasmic factors into the nucleus, maintaining nuclear pore complexes could support innate immunity. Then again, restructuring of these complexes could also benefit viruses that rely on the nucleus for genome replication and immune evasion [69, 70]. Establishing how these trends in gene expression translate to cellular outcomes will require knockout or knockdown of factors related to these pathways and subsequent validation.

Beyond this, our prediction of transcription factors at promoter elements further validates PAF1C as an activator at promoter-proximal sites. Downregulated genes were heavily associated with immune response promoter motifs, including NF-κB components like c-Rel and p65/RelA and coregulator c-Jun/AP-1 (Fig. 5B). This may explain why PAF1-dependent genes were often correlated with the immune response through NF-κB, which plays multiple roles in cytokine expression [71–73]. Moreover, these same promoter-associated motifs were far less enriched in our upregulated genes. Since we associate downregulated genes with PAF1-mediated activation, this result suggests PAF1C triggers this immune response via proximal release of RNAPII. As previously stated, the complex has already been connected to activation at proximal sites [8], so our work fits well into this model of PAF1-dependent gene expression.

Our findings for upregulated immune genes in PAF1 KO are more complex, but they still recapitulate a suppressive role for PAF1C. Though there were some genes found to be immune activators, several others were poised to act in immunosuppressive roles (Fig. 2B). Additionally, PAF1 was found to be suppressive for proviral mechanisms often subject to viral hijacking (Fig. 3C). As such, loss or antagonism of PAF1 may abrogate the immune response by RNAPII release from certain transcription sites. As we had previously shown [14], this could explain why we find positive enrichment for flavivirus host-dependency factors in PAF1 KO (Fig. 4). This effect was diminished for other viruses across weaker immune stimuli. Hence, PAF1 could be required for maintaining a proviral microenvironment for some viruses, but perhaps this only occurs when certain immune pathways are activated during infection.

When we analyzed enhancer motifs associated with our PAF1-dependent genes, our results indicated PAF1C-mediation at these sites may be variable, depending on several factors. For one, a large fraction of enriched enhancer motifs was found to be shared between upregulated and downregulated genes (Fig. 6C). This occurs despite them occupying exclusive genomic regions, suggesting genomic location itself is a stronger determinant of PAF1 functionality at certain sites. Since PAF1C extensively regulates chromatin modification and remodeling [74–78], these roles could be driving such a trend. However, we found a large subset of enhancer motifs exclusively associated with upregulated genes. Several of them were immune-related (Fig. 6), suggesting PAF1 represses certain immune genes through enhancer occupancy. Enhancers associated with upregulated genes also had a tighter genomic distribution and were typically closer to expressed genes compared to those downregulated associated (Fig. S4). This could imply that upregulated genes are regulated by super-enhancers, which are often considered to be motifs in close genetic proximity [79]. Given PAF1C occupancy of super-enhancers has been associated with suppression previously [11], our results support PAF1C restricting gene expression by pausing RNAPII at such sites.

This begs the question as to why we observe enrichment for enhancers associated with downregulated genes and those shared by both gene sets. When we map all motifs to their respective target genes, there is a strong overlap between upregulated and downregulated immune gene targets (Fig. S6). Thus, PAF1C likely induces positive regulation of innate immunity by occupying promoters, in addition to some enhancers. Meanwhile, the complex could further bolster immunity by repressing immunosuppressive genes via enhancer occupancy or activating negative regulators upstream. It suggests a sort of “tug-of-war” between PAF1C acting at proximal versus distal regulatory elements. With this complexity in mind, PAF1C may shift between predominantly active or suppressive roles depending on the context of cell perturbation. This harkens back to our analysis of host-dependency factors, wherein the type of stimuli contributed to the extent of PAF1-mediated suppression. Regardless of the exact mechanisms, our results highlight how models of PAF1C as either an activator or suppressor are not inherently conflicting but may exhibit cross-regulation in the context of innate immunity. It should be noted that PAF1C-mediated suppression of genes has largely been observed in cancer cell lines [9, 11]. Studying if these dual regulatory modes of PAF1C exist in primary immune cells and other epithelial cell lines will be valuable.

These results provide the foundation to further investigate PAF1C occupancy of transcriptional sites in the context of innate immunity. Identifying such sites with ATAC-Seq [80] and ChIP-Seq [81] in PAF1 KO cells would reveal whether our predicted motifs are transcriptionally accessible during the immune response. Also, corroborating our results by studying PAF1C overexpression and monitoring for inverted effects would further strengthen the trends identified here. Strikingly, a concurrent study has already validated aspects of our computational predictions, showing PAF1 to be supportive of NF-κB-dependent gene expression in a siRNA knockdown model [16]. Such work reiterates how our contribution is a critical starting point for future studies focusing on PAF1C and the immune response.

## Conclusions

In summary, we clarify the extent of PAF1C-mediated gene expression and identify notable trends that reaffirm its roles in both activation and suppression of immune gene expression. We show strong correlations between gene expression and known transcriptional motifs. This provides basis for future elucidation of molecular and cellular mechanisms underlying PAF1C function during an immune response. This is especially important since PAF1C can be coopted through host-pathogen interactions to disrupt innate immunity.

## Materials and Methods

### Cells

ncgRNA and PAF1 KO cell lines were generated by CRISPR/Cas9 in A549 lung epithelial cells as previously described [14]. All cell lines were cultured in Dulbecco’s modified Eagle’s medium (DMEM, Gibco ThermoFisher) with 10 % fetal bovine serum (FBS, Gibco ThermoFisher) at 37°C. *Mycoplasma spp.* PCR testing was done monthly.

### Sample preparation

ncgRNA and PAF1 KO cells were plated at a density of 6.0 x 10^5^ cells per well in 6-well plates and incubated at 37°C for 24 hours. Cells were then stimulated with poly I:C (2.0 μg/mL, Tocris), poly dA:dT (1.0 μg/mL, InvivoGen), LPS (1.0 μg/mL, eBioscience), IFN-β (50 U/mL, R&D Systems), and TNF-α (100 ng/mL, Gibco Thermofisher) at the given concentrations. For poly I:C and poly dA:dT, transfection was performed with Lipofectamine 2000 (ThermoFisher) per manufacturer instruction. After 3 hours post-stimulation, all samples were collected with Trizol reagent (Ambion) and stored at −80°C. Total cellular RNA was extracted and purified with the RNeasy mini kit (Qiagen). At the sequencing facility, RNA samples were prepared with the QuantSeq3’ mRNA-Seq Library kit for Illumina (Lexogen GmbH), following a standard protocol [82]. The resulting cDNA libraries were sequenced on an Illumina NextSeq500 system with approximately 4 million reads per sample.

### Quantitative reverse transcription-PCR (qRT-PCR)

Following RNA extraction, sample cDNA was synthesized with the iScript kit (BioRad). Quantitative real time PCR was performed with iTaq SYBR Green premix (BioRad) in 96-well plate format, with data collection and analysis conducted on a LightCycler 480 (Roche). All Ct values were normalized against GAPDH expression. Quantification of gene expression was calculated by the ΔΔCt method, as previously described [83]. Primer sequences are provided in Table S4.

### Immunofluorescence microscopy

Parental A549 cells were fixed with 4% paraformaldehyde following 3-hour treatment with poly I:C (2.0 μg/mL, Tocris), LPS (1.0 μg/mL, eBioscience), and IFN-β (50 U/mL, R&D Systems) at the given concentrations. Cells were permeabilized with 0.25% Triton X-100 and subsequently blocked with 5% goat serum in PBS. Samples were incubated in primary and secondary antibodies for 1 hour each at 25°C. Cell nuclei were stained with Hoechst (Invitrogen). PAF1 primary antibody was diluted at 1:500 (Atlas antibodies, HPA043637). AlexaFluor555 and Hoechst33342, both at 1:1000 dilution, detected PAF1 and Hoechst, respectively. Epifluorescence microscopy images were produced with a Nikon ECLIPSE Ti2 Series microscope using 40x objective. StarDist (Weigert 2018) was used to detect nuclei of cells for measurement of PAF1 fluorescent intensity. All other measurements were performed within Fiji interface (Schindelin 2012).

### Differential gene expression analysis

Read data was evaluated for sequence quality and sample contamination with FastQC (v0.11.9) [84]. Reads were aligned against the human reference genome (build 38) with STAR aligner (v2.5.2b) [85]. FeatureCounts (v2.0.3) calculated read counts for each gene and built a corresponding matrix for all samples [86]. Prior to differential gene expression (DGE) analysis, genes with mean counts less than 1 were excluded. DGE analysis was performed with DESeq2 (v1.34.0) [87], wherein the design accounted for the biological replicate number, type of immune stimuli, and cell type (PAF1 KO, ncgRNA). Direct contrasts between different conditions and cell types were performed after normalization of read counts. Batch effects were removed with ComBat function of the sva package (v3.42.0) [88]. Significant changes in specific genes (p.adj < 0.05) were identified after adjusting for false discovery rate using the Benjamini-Hochberg method.

### Gene set enrichment analysis

Gene-set enrichment analysis (GSEA) was performed with fgsea [89], using Reactome gene sets [90] downloaded from the Molecular Signatures Database [91]. Custom GSEA gene sets for were curated manually from publicly available data and are listed in Table S5. All genes were included in the random walk and ranked by a DESeq2 weighted statistic (stat) for fold change and p-value. Each run performed 10,000 iterations across gene sets, which included sets of 10 to 500 genes for targeted assessment and 50 to 1000 genes for broad assessment. Significant changes in specific gene sets (p.adj < 0.05) were identified after adjusting for false discovery rate using the Benjamini-Hochberg method. Normalized enrichment scores (NES) represent the enrichment score (ES) following normalization to the mean enrichment of random samples of the same gene set size.

### Regulatory elements analysis

The Database for Annotation, Visualization and Integrated Discovery (DAVID) (Knowledgebase v2021q4) [92] was used to assess for statistical enrichment of promoter-associated transcription factor binding sites (TFBS). This was done according to the UCSC Genome Browser database of TFBS conserved tracks, which are experimentally-determined sites [93]. DEGs were inputted and enriched for TFBS with EASE scoring, which produces a modified Fisher’s Exact test p-value [94], correcting for false discovery rate using the Bonferroni method.

Enhancers of DEGs were aggregated with GeneALaCart/GeneHancer as synchronized with GeneCards (v5.7) [95]. Mapping to human genome was done with chromPlot (v1.22.0) [96] and a Jaccard overlap statistic for upregulated-associated versus downregulated-associated enhancers was calculated with the valr package (v0.6.4)[97]. Extracted enhancer regions were filtered as “Enhancer” and “Enhancer/Promoter” and met the criteria of existing in A549 lung epithelial cells and having a GeneCards confidence score > 10 before enrichment. Then, MEME Suite (v4.5.1) program Simple Enrichment Analysis (SEA) [98] identified relatively enriched motifs in the enhancer regions of DEGs. Gene Ontology for Motifs (GOMo) [99] identified which of these enhancer motifs were associated with GO-BP terms. Both MEME Suite programs corrected for false discovery rates using the Benjamini-Hochberg method. Regulatory relationships between transcription factors and target genes were extracted from the Transcriptional Regulatory Relationships Unraveled by Sentence-based Text Mining (TRRUST, v2) database [100].

## Declarations

### Ethics approval and consent to participate

*Not applicable*

### Consent for publication

*Not applicable*

### Availability of data and materials

RNA-sequencing data are available in the ArrayExpress database under accession number E-MTAB-11620. The Sequence Read Archive (SRA) accession number is PRJEB52223. The basic code for the read alignment, differential gene expression analysis, and gene set enrichment analysis is available at: https://github.com/mwkenast/PAF1KO_

## Funding

Funding was provided by the University of California, Davis and the W. M. Keck Foundation.

### Authors’ contributions

MWK and PSS conceived of the work and designed experiments. MWK, OHP, and MJP performed experiments. MWK conducted bioinformatic analyses. MWK and PSS wrote the manuscript. PSS secured funding for work. All authors read and approved the final manuscript.

## Acknowledgements

We would like to thank members of the Shah Lab for constructive criticism and generative discussions. Sequencing was carried out at the DNA Technologies and Expression Analysis Cores at the UC Davis Genome Center, supported by NIH Shared Instrumentation Grant 1S10OD010786-01.

## Supplementary Material

**S1 Figure.**
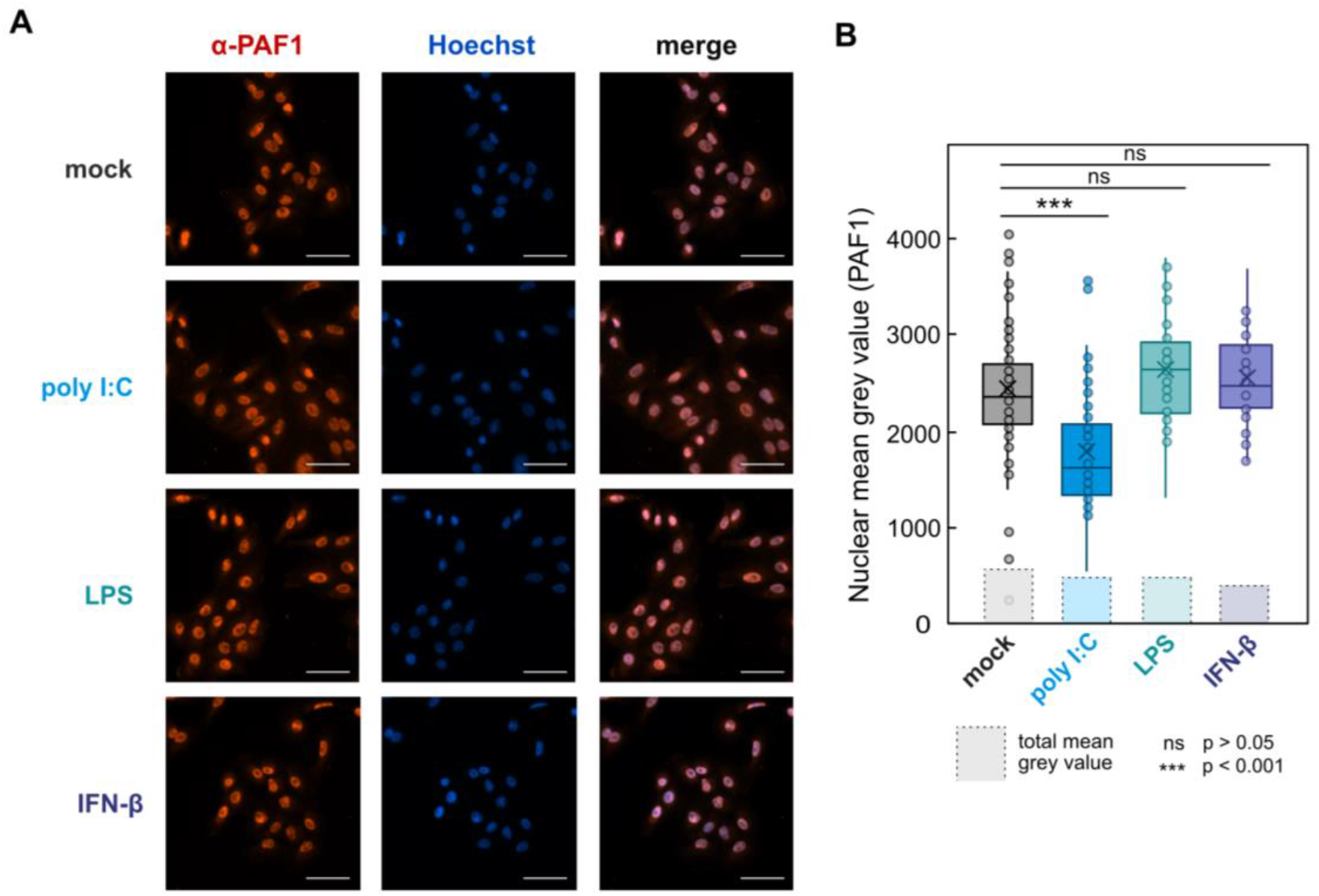
Immune stress marginally affects PAF1 nuclear localization. (A) Immunofluorescence microscopy shows how selected immune stimuli do not significantly disrupt the nuclear localization of PAF1. A549 cells were stimulated for 3 hours with the given stimuli before being fixed and then stained with PAF1 antibody (red) and Hoechst (blue) for marking nuclei. Images were produced with epifluorescence microscopy. All scale bars represent 50 µM. (B) Microscopy images were assessed quantitatively for PAF1 nuclear localization. Cell nuclei were selected for with StarDist, using Hoechst staining as the training control. Grey mean area was used to measure intensity of PAF1 signal in these nuclei (as represented by each point), relative to the average of the total grey mean area for each image. These values were plotted as a distribution. Statistical significance was determined using an unpaired two-tailed Student’s t-test.

**S2 Figure.**
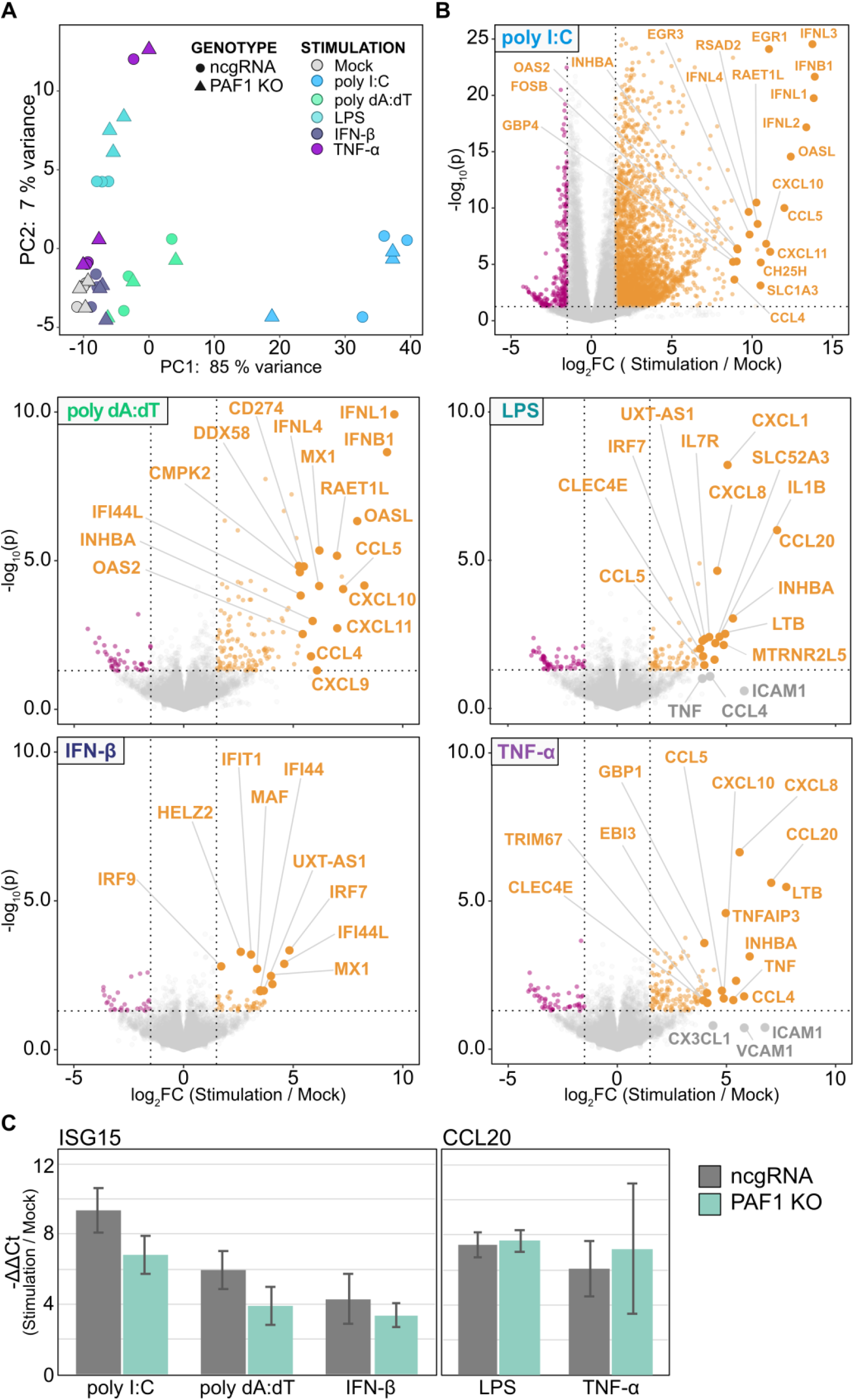
Further characterization of immune activation by stimuli. (A) PCA shows distinct gene expression profile of stimulations relative to mock across both ncgRNA and PAF1 KO cell line. The counts for all genes were corrected with a variance-stabilizing transformation via DESeq2, and only the variability of biological replicates was removed as a batch effect with ComBat. PCA was performed on the resulting counts, with the top 2 principal components being plotted. (B) Volcano plots show activation of the immune response in ncgRNA cell line following each stimulation. Comparisons were performed within the DESeq2 pipeline. The resulting log2 foldchanges and –log10 p-values are plotted for all genes, where top immune response genes are labeled. Cutoffs are set at log2 foldchange > 1.5 or < −1.5, and p-value < 0.05. (C) qRT-PCR analysis validates activation of the immune response in ncgRNA and PAF1 KO cells. Fold changes for ISG15 and CCL20 were calculated using the ΔΔCt method for three technical replicates, being normalized against GAPDH expression. Error bars indicate standard deviation across three biological replicates.

**S3 Figure.**
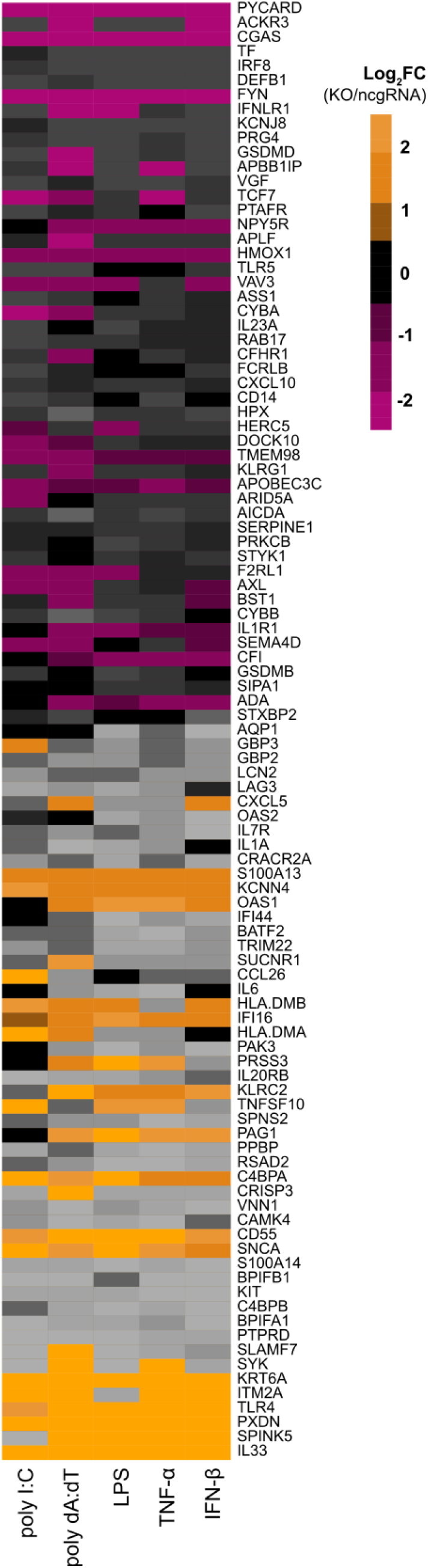
Top PAF1-dependent genes across stimuli. Heatmap shows relative changes in gene expression as log2 foldchanges of DEGs from PAF1 KO versus ncgRNA contrasts of Figure 2. Colors are desaturated for genes with an adjusted p-value > 0.05.

**S4 Figure.**
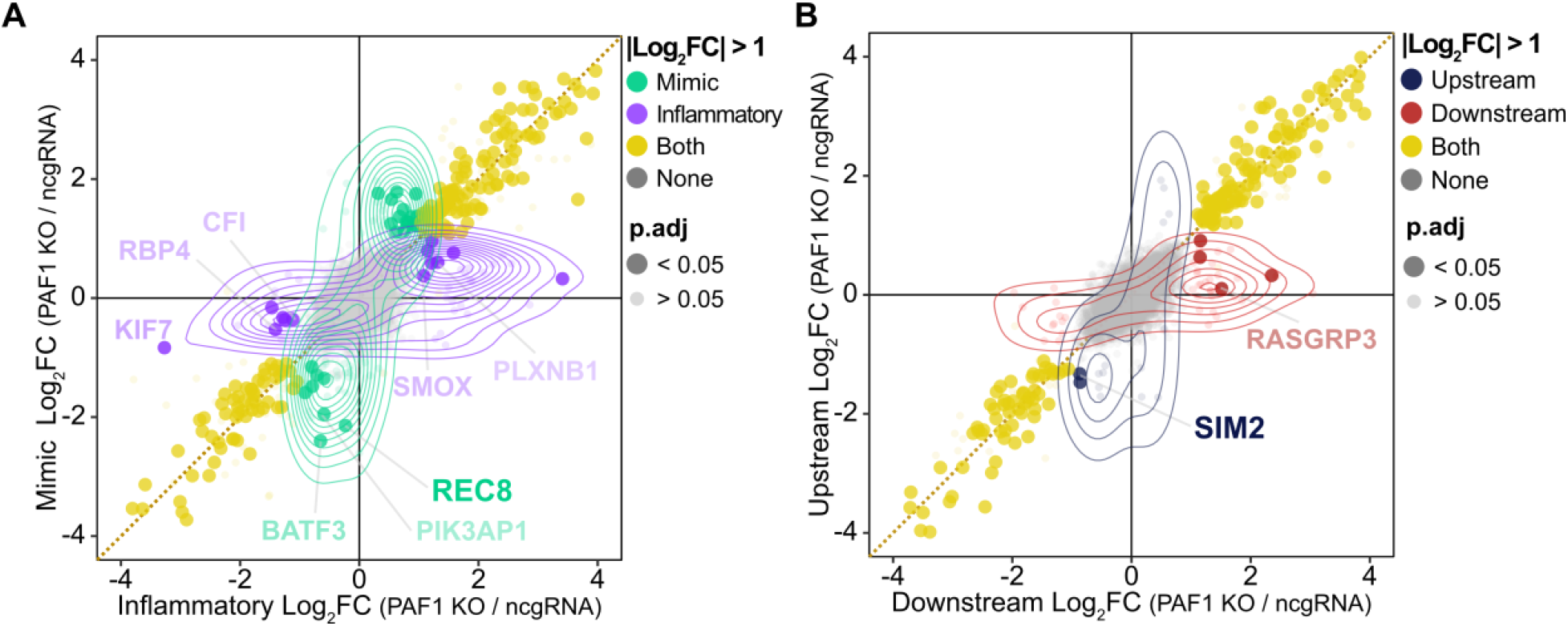
PAF1 alters gene expression unique to some immune stimuli. Dot plots show PAF1-dependent genes include variability across the type of stimulation. (A) The log2 foldchanges and adjusted p-values for PAF1 KO versus ncgRNA cells were averaged across interferon (poly I:C, poly dA:dT) and inflammatory (LPS, TNF-α) stimuli. The average log2 foldchanges were plotted on two axes. Density plots are shown for all genes with log2 foldchanges greater than 1 or less than −1. Significant genes (p.adj < 0.05) are plotted as larger, bolded points and notable genes are labeled. Genes with strong induction (avg. log2 fold-change > 1) in ncgRNA cells are bolded. (B) The same analytical approach discussed above is applied to PAF1 KO DEGs for upstream (poly I:C, poly dA:dT, LPS) versus downstream (TNF-α, IFN-β) stimuli.

**S5 Figure.**
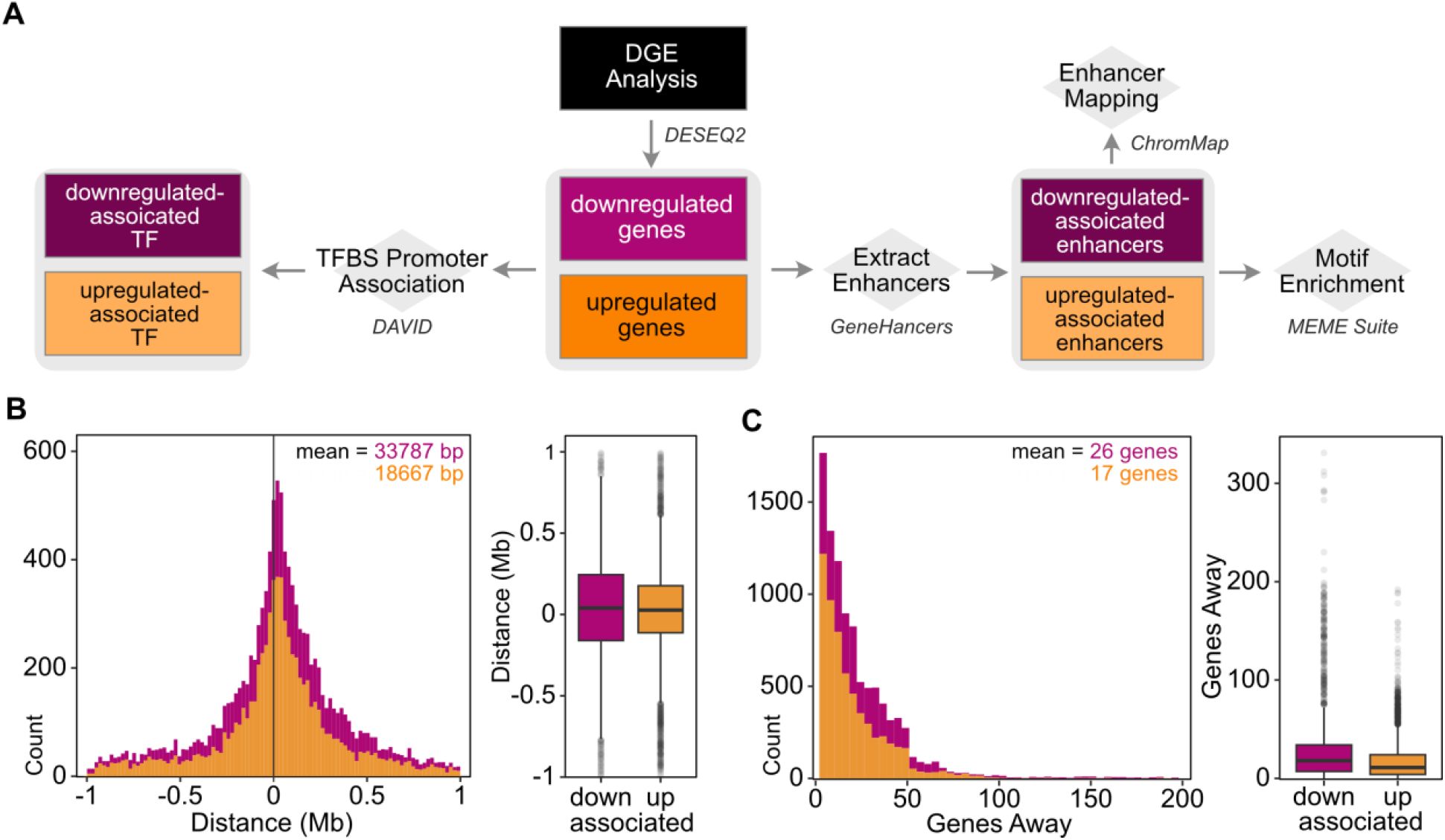
PAF1-associated enhancer analysis pipeline and quality control. (A) Summarization of pipeline for promoter and enhancer motif analyses. (B) Distribution of distances of enhancer motifs from target upregulated (orange) or downregulated (magenta) gene. (C) Distribution of the number of genes in between enhancer motif and target upregulated (orange) or downregulated (magenta) gene.

**S6 Figure.**
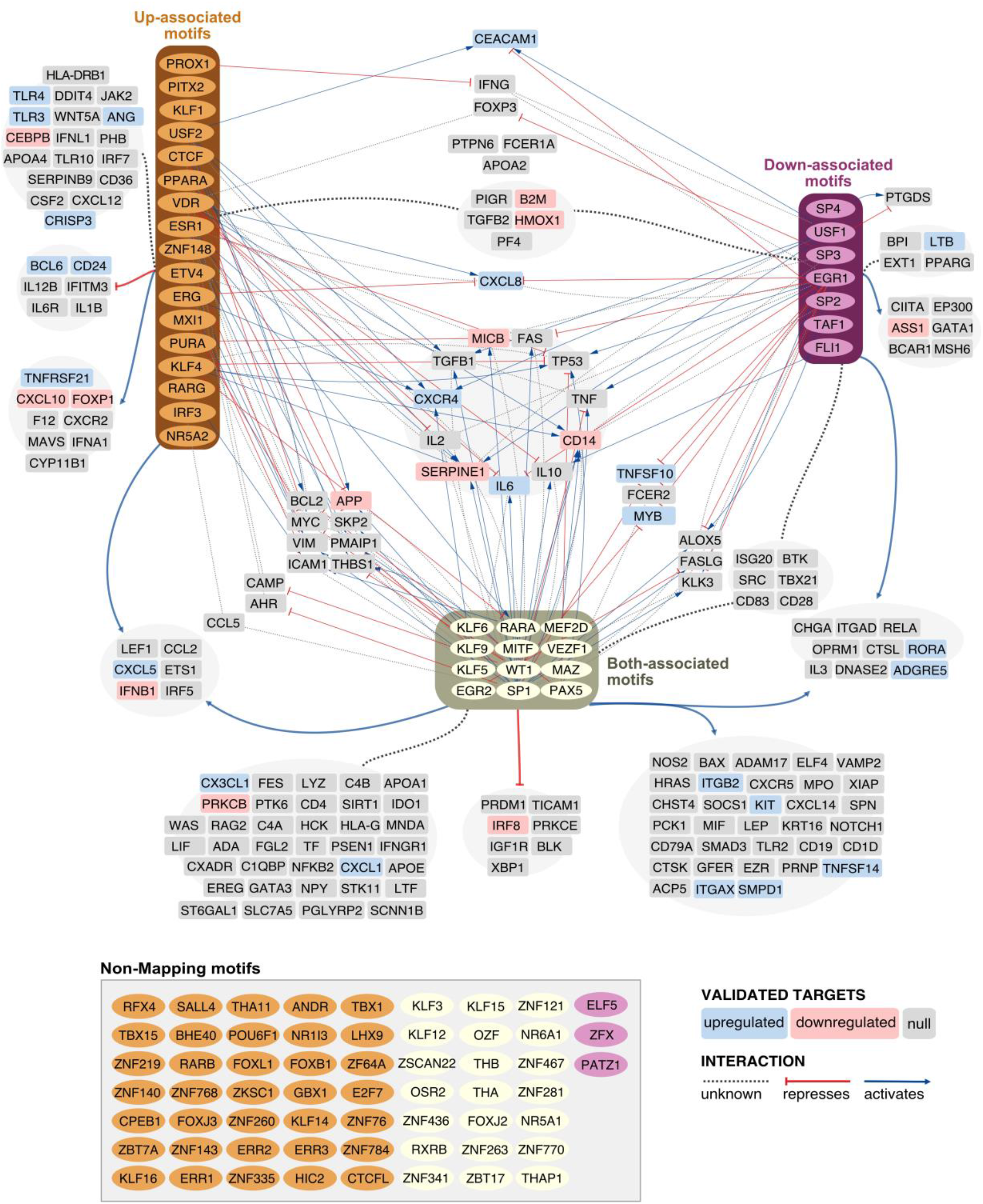
Transcription factor predicted targets network. Identified enhancer motifs associated with known transcription factors map to immune genes (GO:0002376). Significantly enriched enhancer motifs (E < 0.05, enrichment ratio > 2) were screened against the TRRUST v2 database for target genes and the type of interaction. Non-mapping motifs include those that were either not transcription factors or had no known associations in the database. Target genes with consistent differential expression across stimuli for PAF1 KO versus ncgRNA (average log2 foldchange > 1 or < −1) were colored accordingly (red = downregulated, blue = upregulated, grey = did not meet criteria or insufficient reads).

**Table S1. DESeq2 outputs.**

**Table S2. GSEA outputs.**

**Table S3. Associated Promoter and Enhancer Motifs.**

**Table S4. qRT-PCR primers.**

**Table S5. Host-dependency factors.**

